# RNA-SpecFlow: RNA-Conditioned Spectral Flow for Histology Synthesis

**DOI:** 10.64898/2026.10.26.761914

**Authors:** Yiyao Liu, Wenxiao He

**Affiliations:** Glasgow College, UESTC (Univ. Electronic Sci. & Tech. China), Chengdu, China; Faculty of Science, The University of Hong Kong, Pokfulam, Hong Kong

**Keywords:** RNA-conditioned generation, histopathology, flow matching, scale-wise autoregression, spectral fusion

## Abstract

RNA-conditioned histology synthesis models the range of tissue appearances compatible with a transcriptomic representation. We present RNA-SpecFlow, a continuous-latent FlowAR architecture that applies molecular conditioning at adjacent-scale feature transitions. A Biological Latent Adapter maps RNA representations into the generator’s condition space. A shared spectral fusion module separates aligned features into complementary low- and high-frequency bands, modulates each band independently with RNA, and combines them through residual fusion. The generator uses five latent grids of sizes 1, 2, 4, 8, and 16, with per-scale velocity flow matching and an auxiliary objective for retaining the condition in backbone features. We accompany this architectural specification with an analysis of archived outputs from five TCGA cancer cohorts, including 500 saved real/generated tile pairs. Nuclear analysis yields a concrete measurement finding: increased detection counts can coincide with reduced segmentation agreement. On 12 public images with 303 annotated nuclei, twofold enlargement increases HoVer-Net predictions from 287 to 415 while mean panoptic quality changes from 0.673 to 0.458. Switching analysis pipelines also reverses the direction of real/generated count differences in two cohorts. RNA-SpecFlow provides a design for repeated frequency-specific molecular conditioning, and the accompanying experiments establish the sensitivity of morphology-based evaluation to detector inputs and analysis choices.

## 1 Introduction

RNA-conditioned histology synthesis asks what tissue appearances are compatible with a molecular representation. Histology resolves local morphology, while bulk RNA summarizes activity across a heterogeneous specimen. A useful formulation therefore models a one-to-many distribution over tissue regions. Image-to-expression models such as HE2RNA and spatial-transcriptomic predictors demonstrate relationships between morphology and gene expression [1–3]. Generating images in the reverse direction provides a setting for studying how molecular conditions enter a model of tissue appearance.

RNA-CDM established transcriptome-conditioned histology synthesis with cascaded diffusion models [4]. Scale-wise generation introduces a further design question: how should RNA condition the feature transitions between successive resolutions? VAR models next-scale dependencies, and FlowAR combines this hierarchy with continuous-latent flow matching [5, 6]. These transitions provide repeated points at which molecular information can affect feature construction.

We propose RNA-SpecFlow, which applies independent RNA modulation to complementary spatial-frequency bands at adjacent-scale interfaces. The module aligns neighboring features, separates their low- and high-frequency components, and fuses the modulated branches through a residual path. This design gives RNA separate parameters for slowly and rapidly varying latent patterns. A Biological Latent Adapter supplies the shared molecular embedding, and an auxiliary feature-consistency objective encourages its retention in the backbone. The architecture builds on FlowAR’s hierarchy and velocity flow-matching objective; its proposed mechanism is repeated, band-specific conditioning.

Evaluating this design also requires understanding the measurements used to assess generated tissue. Nuclear contours and predicted class proportions offer a local view of morphology, but their interpretation depends on detector calibration. We analyze saved outputs from lung adenocarcinoma (LUAD), glioblastoma (GBM), kidney renal papillary cell carcinoma (KIRP), cervical squamous cell carcinoma and endocervical adenocarcinoma (CESC), and colon adenocarcinoma (COAD). The study combines archived image-distribution scores with nuclear analysis of 500 saved real/generated tile pairs and calibration on 12 public images containing 303 annotated nuclei.

The calibration identifies a mismatch between detection count and instance agreement: twofold enlargement increases HoVer-Net predictions from 287 to 415 while mean panoptic quality changes from 0.673 to 0.458. On saved CESC and LUAD pairs, changing the analysis pipeline reverses the direction of real/generated count differences.

Our contributions are the specification of repeated RNA-conditioned spectral fusion within a five-scale flow generator and an empirical analysis of how measurement choices affect its archived-output diagnostics. The specification covers feature fusion, loss reduction, and the implemented sampling interface. The evaluation retains zero detections, undefined compositions, and complete-pair coverage.

## 2 Related work

### Histopathology generation

PathologyGAN explored adversarial synthesis and latent organization of cancer tissue [7]. Diffusion modeling [8] later supported pathology-specific conditioning and multi-resolution synthesis, including gigapixel whole-slide generation [9]. RNA-SpecFlow addresses local tiles conditioned on bulk transcriptomic representations. Evaluation of synthetic pathology spans image distributions, downstream tasks, and expert assessment, as studied by Pozzi et al. [10]. Our analysis examines the detector-based morphology component of this broader evaluation problem.

### Conditioned image synthesis

Latent diffusion uses a pretrained image autoencoder as a generation workspace [11]. ControlNet introduces spatial controls into pretrained diffusion models [12]. Pathology-specific conditions include nuclei-aware masks in NASDM and report semantics in PathLDM [13, 14]. Nucleus masks specify locations, while bulk RNA describes a specimen without locating individual cells. RNA-CDM studies the RNA-to-image direction through cascaded diffusion [4]. RNA-SpecFlow investigates repeated frequency-specific conditioning within a scale-wise flow architecture.

### Molecular and morphological learning

HE2RNA predicts RNA expression from whole-slide images [1]; ST-Net and Hist2ST use spatially resolved expression targets to study local morphology and expression [2, 3]. PathGen synthesizes transcriptomic features from histopathology representations for multimodal prediction [15]. These approaches learn from image-to-expression or spatially supervised relationships. Bulk-RNA-conditioned image synthesis instead models a distribution over tissue regions linked to a specimen, making biological pairing and output-level molecular assessment central to its evaluation.

### Scale-wise flow and frequency processing

VAR models next-scale dependencies [5], and FlowAR learns continuous latent groups with per-scale flow matching [6,16]. These mechanisms supply RNA-SpecFlow’s generation backbone. Fast Fourier Convolution and Focal Frequency Loss study feature processing and reconstruction objectives in the frequency domain [17, 18]. Our fusion module uses rectangular Fourier masks and RNA-dependent channel modulation on aligned scale features. MeanFlow learns interval-average velocities through a distinct training identity [19]; RNA-SpecFlow uses instantaneous velocity regression. Section 3 specifies its two-time sampler and the difference between that interface and diagonal-time supervision.

## 3 Methodology

RNA-SpecFlow models a distribution of histology tiles conditioned on RNA. A Biological Latent Adapter supplies molecular embeddings to a scale-wise FlowAR backbone and to a shared spectral module at neighboring resolutions. Velocity flow matching trains the generator, with an auxiliary objective for condition retention in backbone features. This section specifies the RNA-only architecture and its implemented training and sampling interfaces.

### 3.1 Problem formulation and frozen latent interface

Let 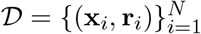, with H&E tile **x**_i_ ∈ ℝ^3×256×256^ and RNA representation 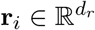. The target distribution *p*_θ_(**x** | **r**) describes the range of tissue regions associated with a bulk RNA profile through specimen linkage and tile selection. Each profile can correspond to multiple local appearances.

A frozen image encoder *E*_A_ and decoder *D*_A_ provide the intended VQ-VAE latent interface:

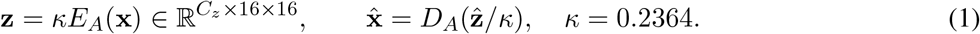

Here *E*_A_ returns continuous latents, and inverse scaling follows the evaluation code. The training engine calls `encode(x).sample`(), which requires a posterior-sampling interface. *C*_z_ denotes the latent channel count. The generator predicts continuous latents.

### 3.2 Overview of RNA-SpecFlow

Figure 1 shows the RNA-only architecture. Five latent groups supply targets at resolutions 1, 2, 4, 8, 16. The adapter maps RNA to the backbone width. Shared encoder–decoder Transformer blocks produce current-scale features from earlier-scale context; a shared spectral module operates at the four adjacent-scale transitions. A conditional velocity field generates continuous latent tokens at each scale. The auxiliary consistency objective uses unfused backbone decoder features. During inference, generated groups replace teacher-forced context, and the final 16 × 16 group is decoded into an image.

**Figure 1.**
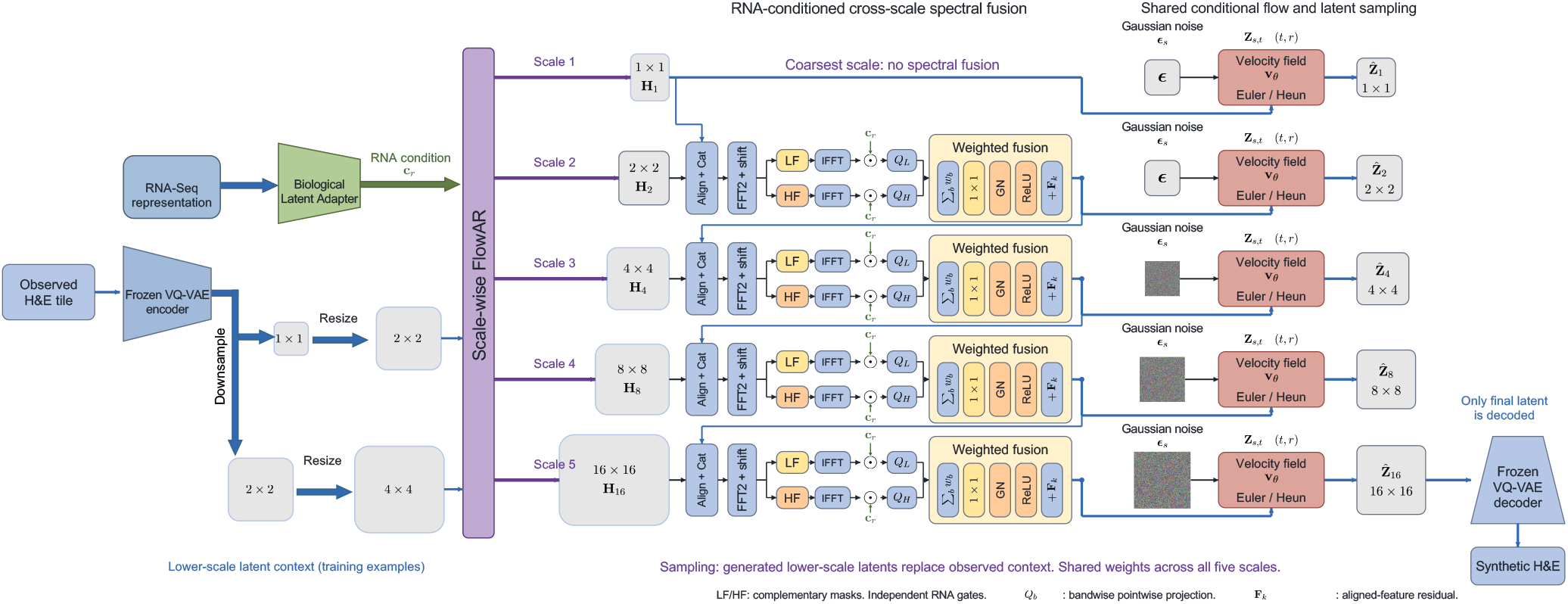
RNA-SpecFlow architecture. A shared FlowAR backbone produces five scale features. The first scale has no spectral fusion; each subsequent module combines the previous fused feature with the current backbone feature. “Align + Cat” abbreviates spatial processing, alignment, concatenation, and normalization. LF/HF denote complementary frequency masks, and IFFT denotes inverse shifting and transformation. Green **c**_*r*_ ports represent independent bandwise RNA gains 1 + ***γ***_*b*_, followed by projections *Q*_*b*_ and weighted residual fusion with **F**_*k*_. Each pink block combines a shared conditional velocity field with multi-step Euler/Heun-style sampling from Gaussian noise; only the final generated 16 × 16 latent is decoded. The left-hand latent grids illustrate lower-scale training context, which is replaced by generated context during sampling. All repeated backbone, fusion, and velocity blocks share parameters. Losses and the biological consistency head are described in the text; Figure 2 expands the fusion module.

### 3.3 Multi-Scale latent autoregressive modeling

With 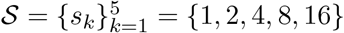, area resizing and tokenization construct

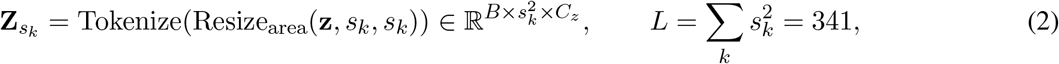

where *B* is batch size and *L* the total token count. Following next-scale modeling in VAR and FlowAR [5, 6], the intended factorization is

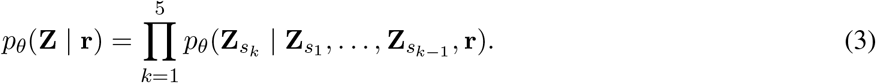

Attention is bidirectional within each scale, and a scale-causal mask blocks finer future groups.

Shifted inputs exclude clean current-scale targets from the corresponding feature. The first-scale input is the RNA embedding. For *k >* 1, training projects the previous target group after bicubic resizing to 16 × 16 and area resizing to *s*_k_ × *s*_k_. The input at group *k* therefore comes from 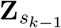. A shared RNA token precedes the projected groups. The decoder outputs 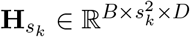, where *D* is backbone width. Sampling directly bicubic-upsamples the generated previous group and caches earlier attention keys and values.

### 3.4 Biological latent adapter

The adapter maps the input representation to width *D*:

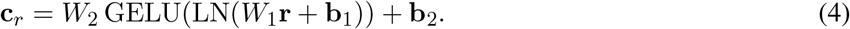

Layer normalization and nonlinear projection map RNA features into the image-latent condition space. The embedding supplies FlowAR’s initial token and adaptive-normalization condition, as well as the molecular input to spectral fusion. RNA reaches the velocity network through the resulting scale features.

Condition dropout replaces the embedding with a learned unconditional vector:

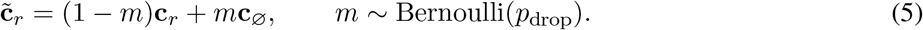

The backbone and fusion module receive the same dropped condition, training both conditional and unconditional feature paths. The inference routing is specified below.

### 3.5 RNA-Conditioned cross-scale spectral fusion

Let 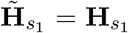. For *k >* 1, the shared fusion module consumes the *previous fused feature* and current unfused feature:

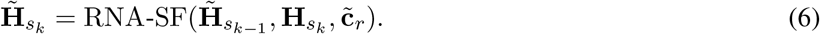

Token groups are reshaped to *B* × *D* × *s* × *s* grids. Figure 2 shows the module’s three operations.

**Figure 2.**
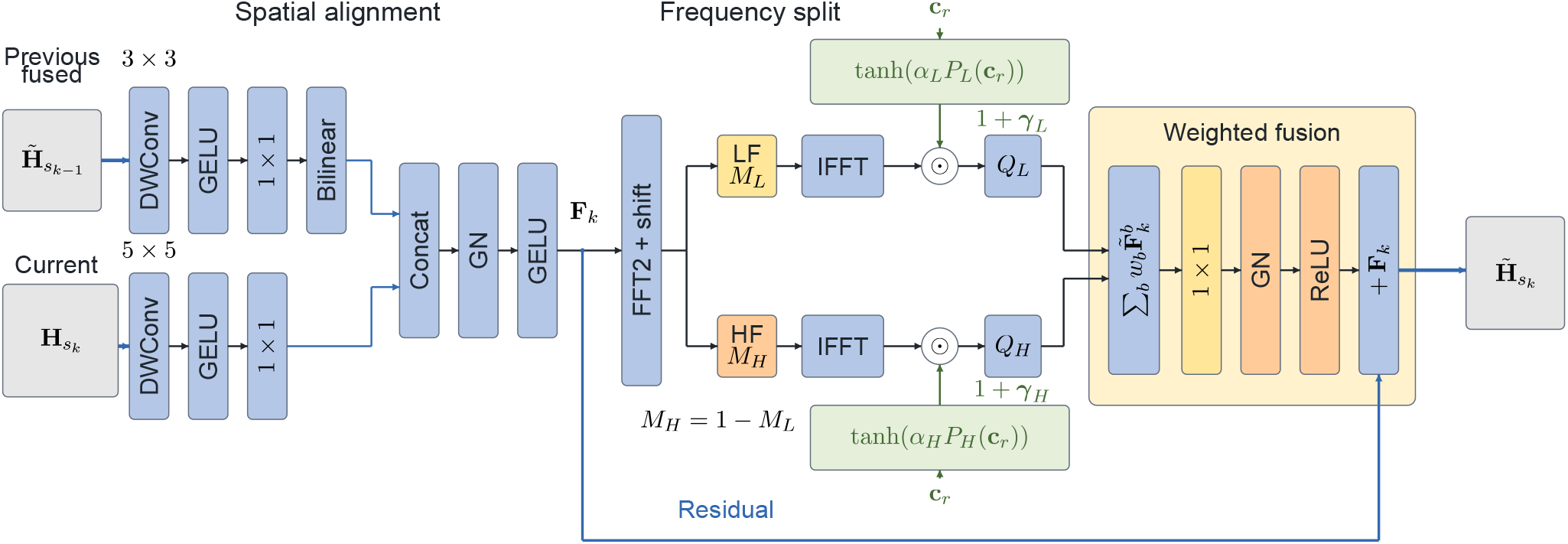
RNA-conditioned cross-scale spectral fusion. The previous fused feature and current FlowAR feature are spatially processed and aligned, split into complementary Fourier bands, and separately modulated by the shared RNA embedding. Learned band weights and output projection are followed by a residual addition of the aligned feature **F**_*k*_. The two frequency branches have independent modulation parameters; the fusion module is shared across the four scale transitions.

#### Spatial alignment

The previous feature passes through a 3 × 3 depthwise convolution, GELU, and a 1 × 1 projection to *D/*2 channels; the current feature uses the same sequence with a 5 × 5 kernel. Bilinear interpolation aligns the previous feature to the current grid. Concatenation restores *D* channels:

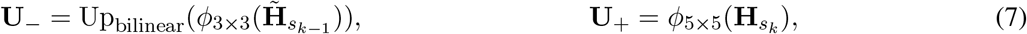

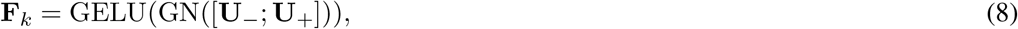

where *ϕ* includes the depthwise convolution and projection, [·; ·] concatenates channels, and GN is group normalization. Different kernels introduce scale-dependent spatial processing before spectral decomposition.

#### Frequency decomposition

Frequency-domain processing separates spatial variation in the learned feature grids [17, 18]:

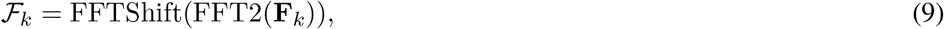

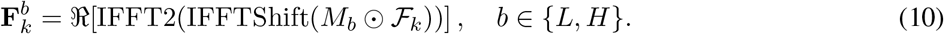

The low-pass mask *M*_L_ is a centered rectangle; *M*_H_ = 1 − *M*_L_ is its complement. In each dimension of length *s*_k_, the integer radius is ⌊0.5⌊*s*_k_*/*2⌋⌋, with inclusive endpoints clipped to the grid. The mask-construction parameter is 0.5; the four fusion grids retain 1, 9, 25, and 81 low-pass coefficients per channel, respectively.

#### RNA-aware modulation and residual fusion

Each band uses its own Linear–LayerNorm–ReLU projection *P*_b_, scalar *α*_b_ initialized to 10^−5^, and 1 × 1 projection *Q*_b_:

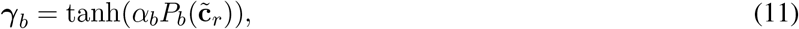

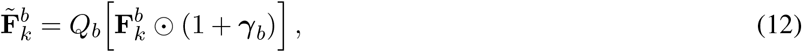

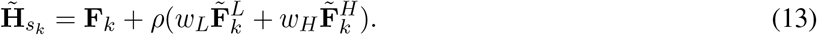

Modulation broadcasts over spatial positions. The scalar weights *w*_L_, *w*_H_ are learned independently and start at one; *ρ* is a 1 × 1 projection followed by GN and ReLU. Near-zero *α*_b_ initializes the molecular gains close to one, while the remaining fusion operations retain their learned transformations. This construction applies RNA-dependent gains at every adjacent-scale transition, with separate parameters for each frequency band.

### 3.6 Conditional velocity flow matching

For target **Z**_s_, sample *t* ∼ *U*(0, 1) and ***ϵ***_s_ ∼ N (0, *I*) independently per example and scale. The linear path runs from data at *t* = 0 to noise at *t* = 1:

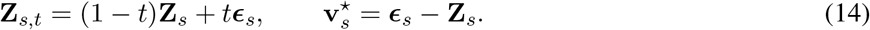

The Transformer velocity field uses adaptive normalization and the scale feature 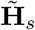. Its two time arguments coincide during training:

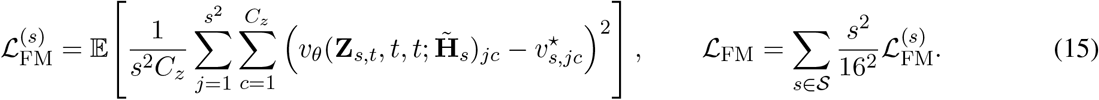

The expectation includes the batch average. Each scale averages over tokens and channels, matching SILoss. Area weighting then gives equal aggregate weight per token across scales under equal errors. The objective regresses instantaneous velocity [16]; MeanFlow’s interval-velocity identity and derivative-based target define a different objective [19].

### 3.7 Biological consistency regularization

Unfused backbone decoder tokens are pooled before spectral processing:

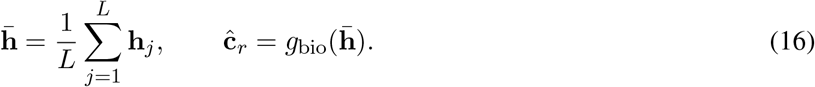

#### Algorithm 1 RNA-SpecFlow training and scale-wise sampling

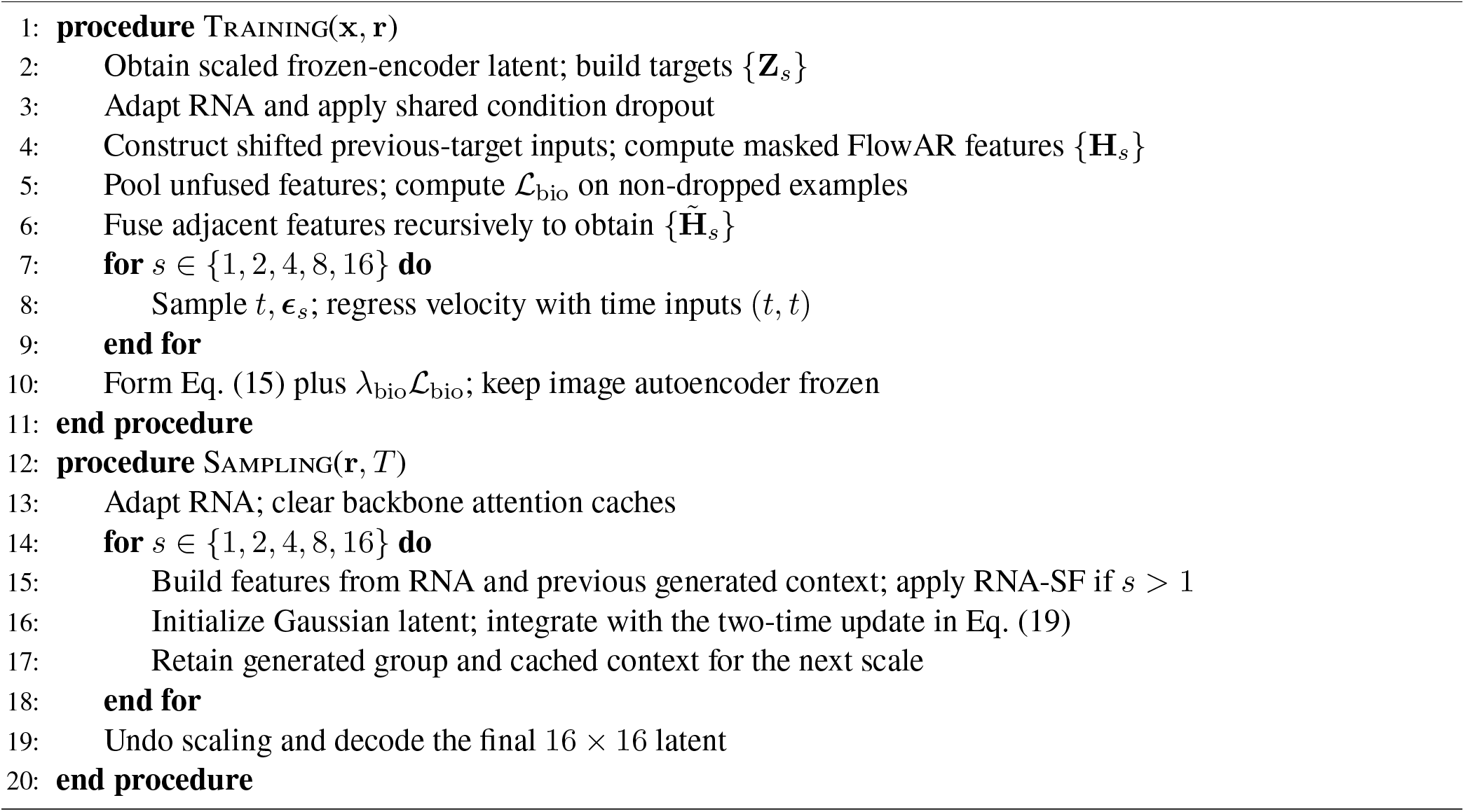

For the set *ℐ* of examples whose RNA condition is not dropped, the auxiliary loss is

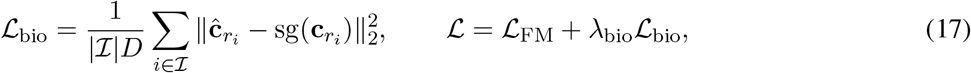

The loss is zero when *ℐ* is empty. Stop-gradient blocks the target branch; gradients still reach the adapter through the conditioned backbone. The objective measures feature retention.

### 3.8 Training and coarse-to-fine sampling

Algorithm 1 summarizes the data flow. At inference, each scale starts from Gaussian noise and is integrated from 1 to 0 before its latent supplies the next scale. Attention caches retain already processed context. For consecutive times *t*_i_ *> t*_i+1_ and Δ_i_ = *t*_i+1_ − *t*_i_, the available sampler evaluates

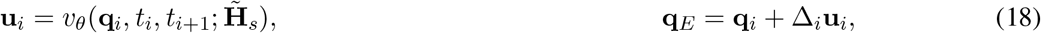

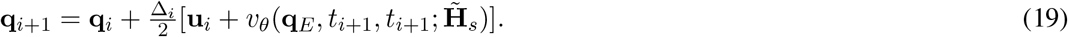

The final interval uses the Euler proposal. With *T* intervals, Heun correction requires 2*T* − 1 field calls per scale; Euler requires *T* .

Classifier-free guidance is intended to use 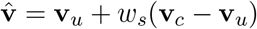 with a scale-ramped *w*_s_.

### 3.9 Implementation scope

RNA-SpecFlow uses the large backbone with width 1024, eight encoder and eight decoder blocks, and 16 attention heads. RNA input width is 768; the velocity predictor has depth 12 and width 1536. Configuration values are *p*_drop_ = 0.1, *λ*_bio_ = 0.5, and 25 sampling intervals. The RNA-only configuration sets use_topo=False, selected through –no_topo in the entry points.

## 4 Preliminary evaluation

We examine image appearance across five cancer cohorts and test how nuclear-analysis choices affect the interpretation of saved real/generated pairs. Archived distributional scores describe image features; annotated controls and pipeline perturbations measure detector behavior. Table 1 defines the evaluation populations.

**Table 1.** Evaluation populations and analysis units. The generation budget is filename-reported; tile counts refer to the saved image archive.

| Evidence | Observed scope | Supported use |
| --- | --- | --- |
| Metric JSONs | Six metric summaries | Archival image-distribution scores |
| Saved JPEGs | 100 pairs/cohort; six cohorts | Traceable appearance panels |
| Nucleus analysis | 1,000 tiles; five cancer cohorts | Fixed-analyzer counts and predicted proportions |
| Paired composition | 365 of 500 pairs with classified nuclei on both sides | Conditional, tile-weighted analyzer differences |
| Public nuclear calibration | 12 images; 303 published annotated nuclei | Input/pipeline calibration |

### 4.1 Artifacts and evaluation provenance

Six JSON summaries record FID, Inception Score (IS), Precision, and Recall for five cancer cohorts and pooled TCGA-ALL. Filenames specify a nominal 50,000-image budget, 50 sampling intervals, and a last-checkpoint label; each JSON records requested CFG 3.5. We report the recorded values as archival point estimates and exclude pooled TCGA-ALL from equal-cohort macro averages.

The evaluation code uses the OpenAI diffusion evaluator’s TensorFlow Inception-v3 graph. FID compares 2048-dimensional pool_3 features; IS uses softmax predictions in groups of 5,000; Precision/Recall use a three-neighbor manifold estimator. Inputs are image arrays on the 0–255 scale, and the real reference population comes from the evaluation loader.

Each cohort directory contains 100 complete 512 × 256 JPEG composites. The saving code places the real 256 × 256 tile on the left and the generated tile on the right. It saves the first 100 loader-ordered pairs before shuffling generated images for metric computation. Display indices retain the saved real/generated pairing. Appearance and contour panels retain the saved pixels. Transformed inputs are used in separately labeled diagnostic experiments. Excluding pooled ALL gives 500 real and 500 generated tiles for nuclear analysis.

### 4.2 Archived image-distribution scores

Across the five cancer cohorts, equal-cohort averages are FID 18.549, IS 2.153, Precision 0.741, and Recall 0.361 (Fig. 3). GBM has the lowest FID and highest Recall within the archive; COAD has the highest FID and lowest Recall. Pooled ALL describes a separate reference mixture.

**Figure 3.**
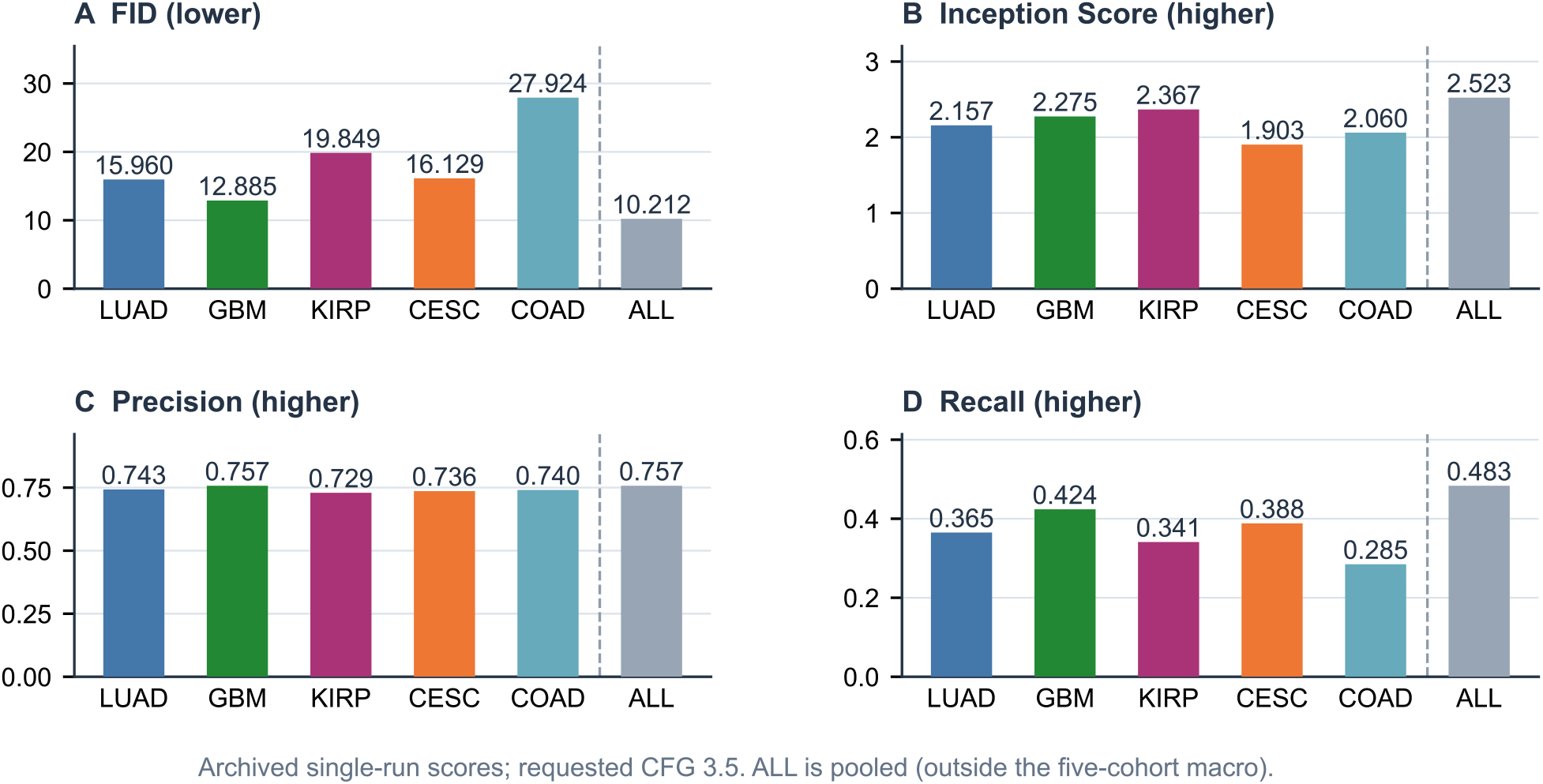
Archived distributional image scores. Values are archived point estimates with requested CFG 3.5 and filename-reported 50 sampling intervals. Dashed separators distinguish pooled ALL from the five cancer cohorts.

### 4.3 Traceable tissue examples

Figure 4 shows two saved pairs per cohort, selected before nuclear analysis. Sample A is nearest the cohort’s multivariate median of generated-side tissue coverage, luminance, saturation, and sharpness. Sample B lies within the proxies’ interquartile ranges and maximizes robust distance from A; an empty eligible set triggers selection of the next-nearest median sample. Each real half retains its saved partner. The panels show variation in tissue color and texture; Appendix A lists the indices.

**Figure 4.**
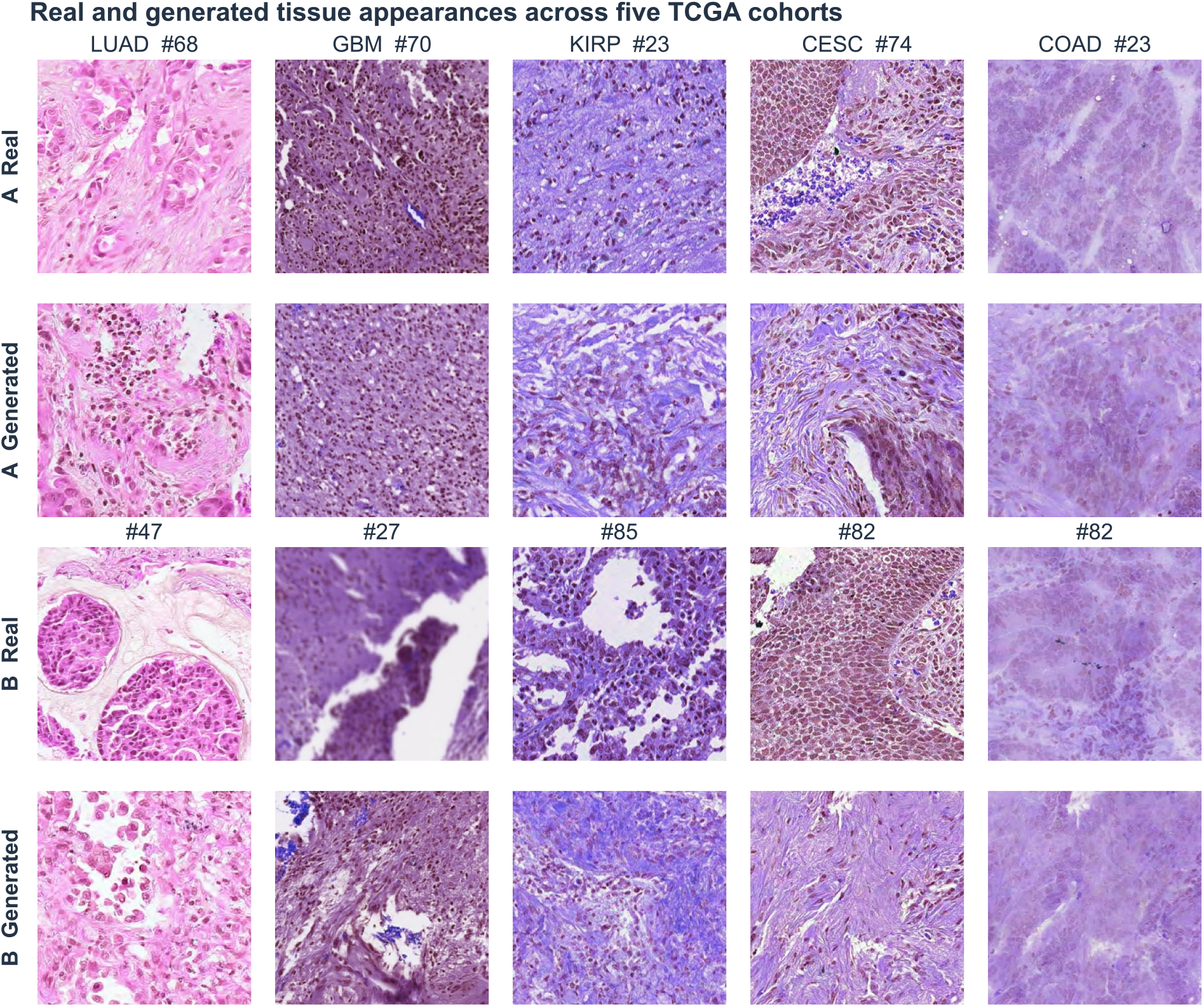
Unmodified real and generated JPEG tile halves from five cohorts. A and B denote two deterministically selected display pairs per cohort; saved indices are shown. Real/generated pairs follow the saved composite layout. The same indices are used in the contour panels, avoiding reselection based on analyzer response. Each tile has 256 × 256 native pixels.

### 4.4 Fixed-Analyzer nuclear morphology

#### Model and resolution assumption

We use TIAToolbox’s hovernet_fast-pannuke model [20–22], verifying the weight file’s SHA-256 before inference. The legacy analysis assumes 0.5 *µ*m/pixel and internally resamples to the model’s 0.25 *µ*m/pixel resolution. Inference uses CPU, batch size 4, zero loader workers, seed 20260904, and disabled automatic tissue masking. Whole-image inference merges overlapping predictions and maps coordinates to native tile pixels.

#### Labels and estimands

We retain argmax instance types and probabilities without additional confidence or morphology filtering. Tumor, Inflammatory, Connective, Dead, and Normal are model-class aliases. Inflammatory includes categories beyond lymphocytes; Normal denotes a predicted nuclear class. Unclassified instances contribute to counts and contour displays but are excluded from class-proportion denominators.

For class *c* and tile *i*, define

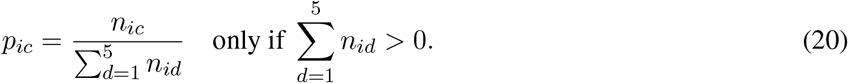

Composition is undefined for tiles with no classified instance. Figure 5A uses each side’s eligible tiles. Paired estimates use *ℰ*_k_, the saved indices with defined composition on both sides in cohort *k*:

**Figure 5.**
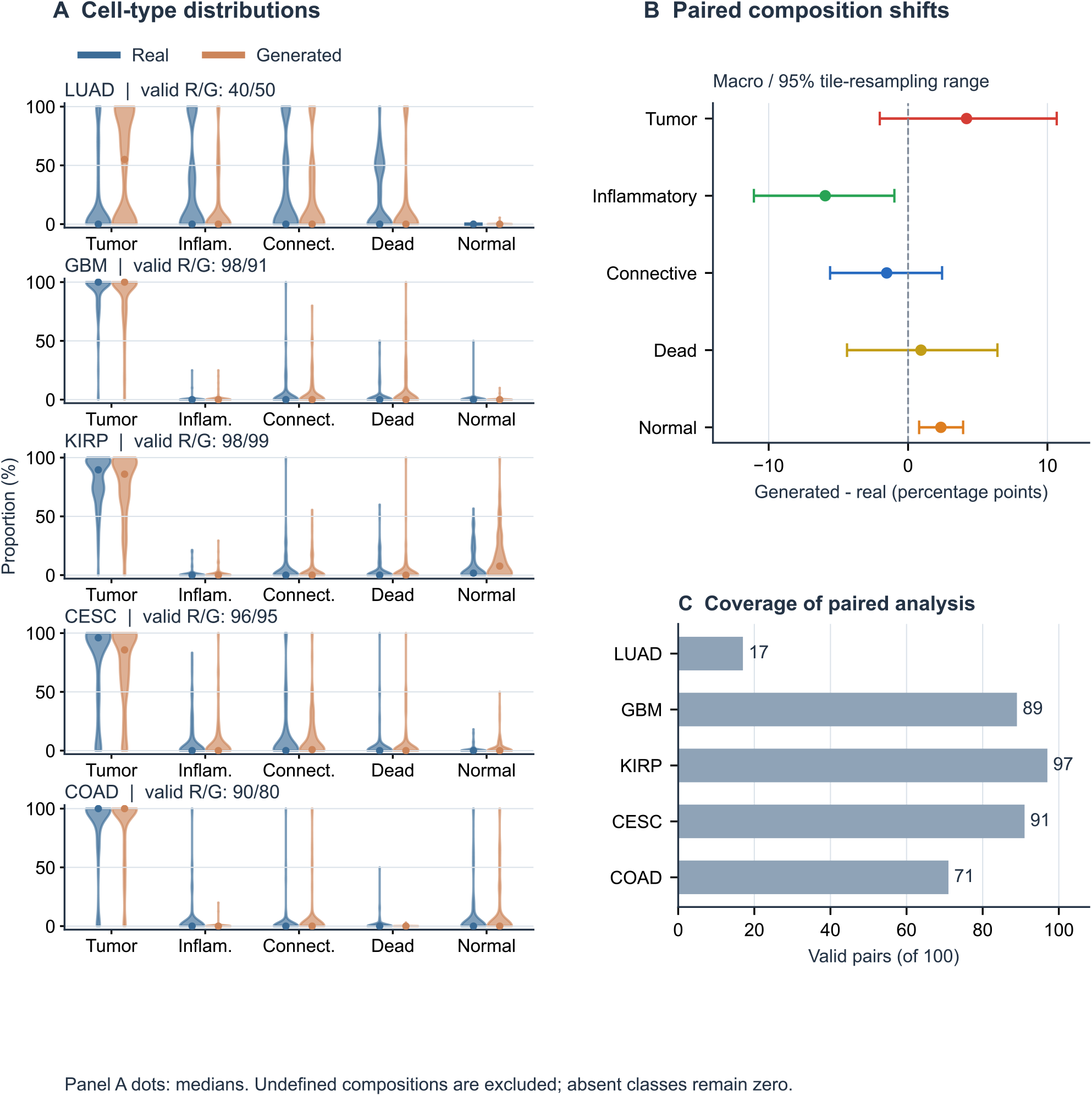
Predicted nuclear composition and analysis coverage. (A) Per-tile distributions on each side’s eligible subset; dots denote medians and R/G labels give valid real/generated tile counts. Violins show kernel-density summaries; zero class fractions are retained and undefined tile compositions are omitted. (B) Equal-cohort paired differences and 95% conditional tile-resampling ranges. (C) Complete-pair counts out of 100 saved indices per cohort. Responses describe model predictions on eligible tiles.

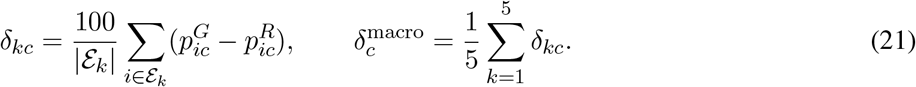

Superscripts *G, R* denote generated and real. Differences are in percentage points and summarize complete pairs; panel A shows side-specific eligible distributions with median dots. We use 10,000 percentile bootstrap resamples of saved pairs within cohorts, retaining undefined entries and recomputing eligible means. Macro resamples weight cohorts equally. The 95% ranges describe conditional tile-resampling variation.

#### Coverage and observed responses

The analyzer predicts 37,073 instances across 1,000 tiles, including 157 zero-instance outputs (75 real, 82 generated). Composition is defined in 422 real and 415 generated tiles. The 365 complete pairs comprise 17, 89, 97, 91, and 71 indices for LUAD, GBM, KIRP, CESC, and COAD. Counts include zeros; Appendix B gives cohort details. LUAD’s paired composition estimate thus describes its 17 eligible pairs.

Equal-cohort paired shifts are +4.19 pp for Tumor, −5.94 pp for Inflammatory, −1.53 pp for Connective, +0.93 pp for Dead, and +2.36 pp for Normal. The displayed ranges cross zero for Tumor, Connective, and Dead.

#### Per-nucleus contour visualization

Figure 6 overlays every saved predicted nuclear boundary on the selected original tiles, colored by predicted class and accompanied by an instance count. The same indices as Fig. 4 retain sparse and zero-instance outputs. These model predictions motivate the annotated calibration and input-sensitivity experiments below.

**Figure 6.**
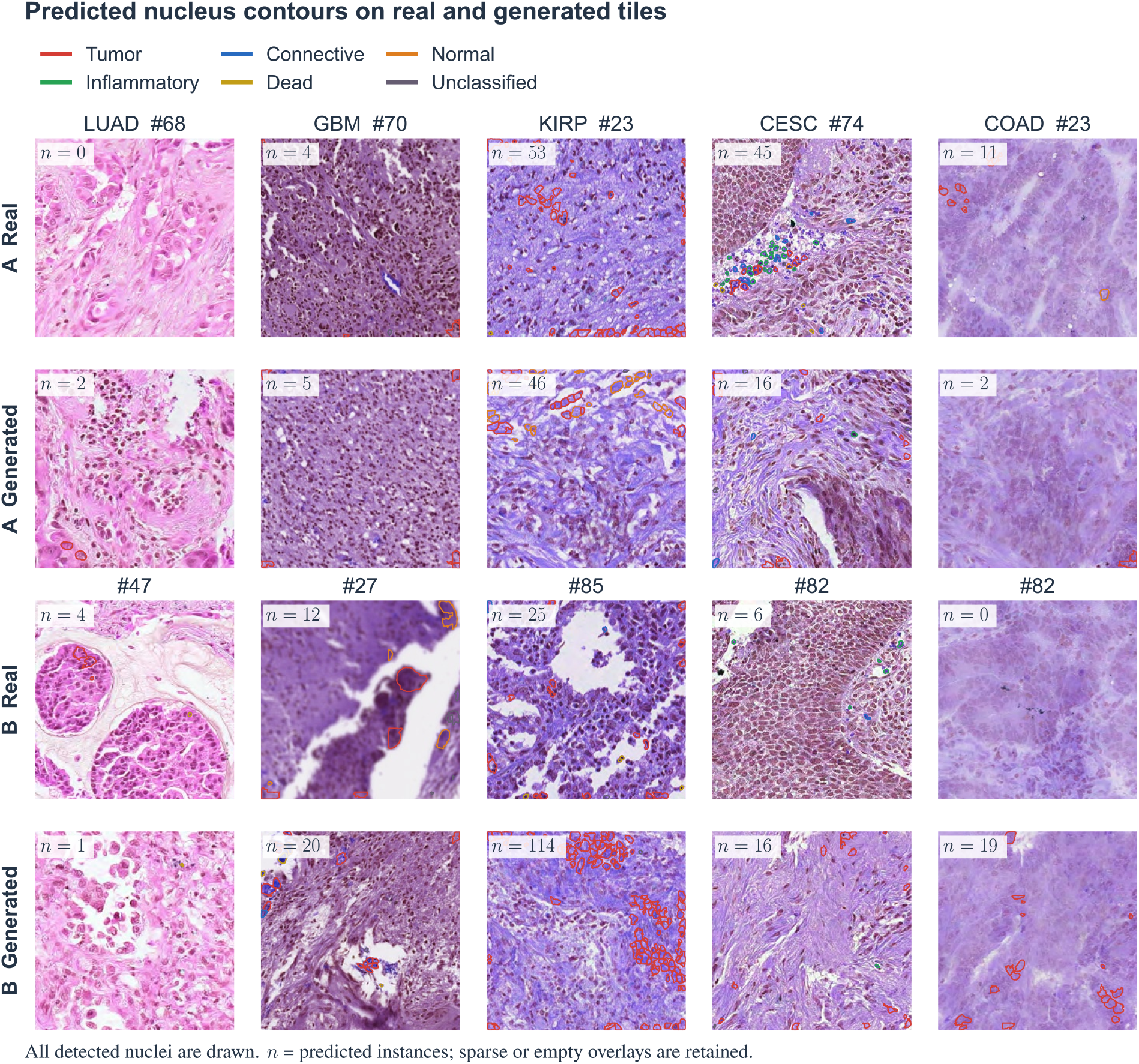
Per-nucleus predicted boundaries on the same A/B tiles as Fig. 4. Colors encode model-assigned classes, including Unclassified; *n* counts all predicted instances in the displayed tile. Every instance from the saved analyzer output is drawn. The original pixels are unchanged. Contours are model predictions, with sparse and zero-instance outputs retained.

#### Resolution sensitivity pilot

Changing the assumed source MPP from 0.5 to 0.25 on the ten fixed Sample A halves reduces the predicted instance total from 184 to 6 and increases empty outputs from one to six. Model weights and inference settings are held fixed. Appendix B gives cohort counts.

### 4.5 Analyzer calibration and measurement sensitivity

#### Published-annotation control

The annotated control contains three images each from Colon, Lung, Kidney, and Cervix in a public PanNuke fold-2 mirror. Selection follows row order and requires at least ten nonoverlapping annotated nuclei per image before inference. One candidate with overlapping masks is excluded and recorded, leaving 12 images and 303 annotated nuclei. PanNuke combines automated labeling and expert validation [21]. This subset measures calibration response; its exact rows and revision appear in Appendix C.

Instances are matched one-to-one at mask IoU *>* 0.5, retaining boundary fragments. Precision and recall pool matched, false-positive, and missed instances across images; panoptic quality (PQ) is averaged over images. Figure 7 compares canonical RGB, twofold pixel enlargement, red/blue channel exchange, and controls for blur, stain normalization, and JPEG encoding.

**Figure 7.**
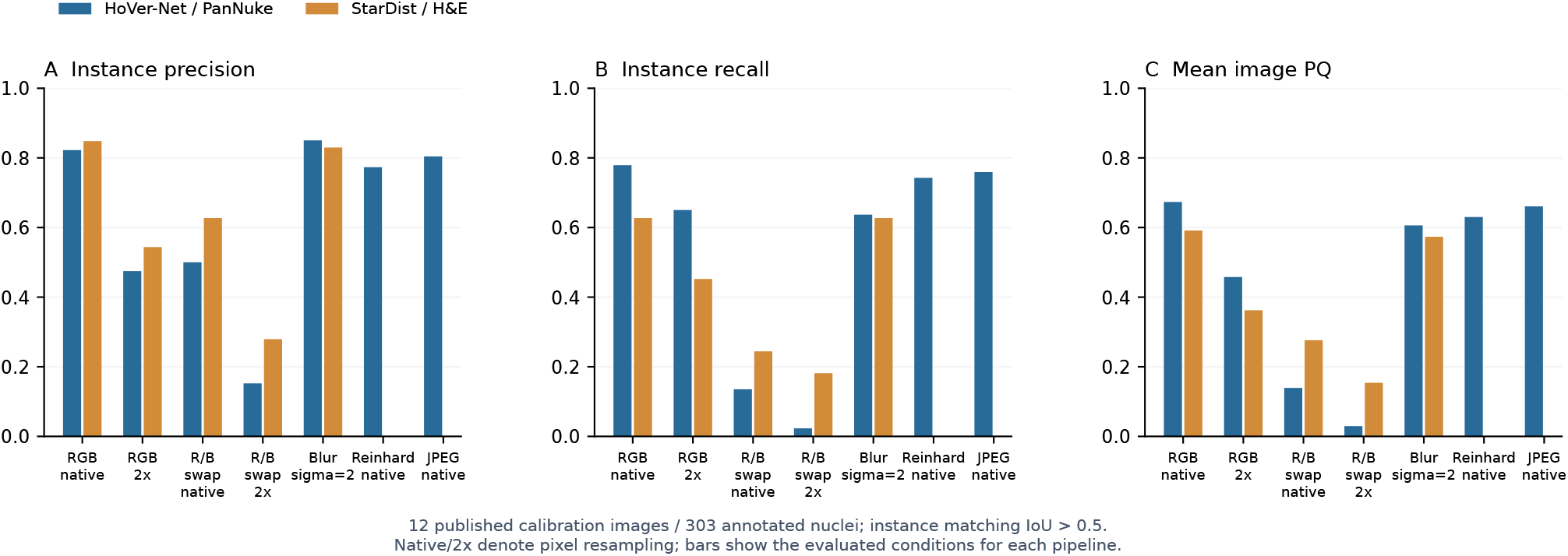
Calibration on 12 public images with 303 annotated nuclei. A/B pool instance matches; C averages image PQ. Both detectors use the same images and published masks. Native and 2× denote pixel resampling. R/B swap exchanges red and blue channels; blur uses Gaussian *σ* = 2; Reinhard uses a fixed reference image; JPEG uses historical quantization tables and chroma subsampling. Absent bars mark untested conditions.

On native RGB inputs, HoVer-Net obtains precision 0.822, recall 0.779, and mean PQ 0.673. Twofold enlargement increases predictions from 287 to 415 while precision changes to 0.475, recall to 0.650, and PQ to 0.458. Native-scale red/blue exchange gives recall 0.135. Detection counts and segmentation agreement therefore respond differently to resampling, and channel order strongly affects recall. StarDist [23], with its released H&E weights, obtains native RGB recall 0.627 and PQ 0.591 on the same control.

#### Historical-image pipeline dependence

We evaluate an independent StarDist pipeline on all 100 saved pairs in each of CESC, COAD, and LUAD, under saved color and R/B exchange with twofold resampling (1,200 tile conditions). StarDist uses the released 2D_versatile_he model, default probability/NMS thresholds, and channelwise percentile normalization. The pipeline comparison varies detector architecture, pretrained weights, and preprocessing together. All zero detections are retained.

For CESC, COAD, and LUAD, respectively, generated/real mean detected-count ratios are 0.617, 0.573, and 1.531 under legacy HoVer-Net, compared with 1.576, 0.932, and 0.837 under saved-color StarDist. R/B-exchanged StarDist gives 1.302, 0.986, and 1.056. Figure 8 shows 95% ranges from 10,000 paired tile resamples (seed 20261007). CESC and LUAD cross the equal-count threshold when the pipeline changes, so the direction of the difference depends on analysis choices. The ranges describe variation across saved tiles.

**Figure 8.**
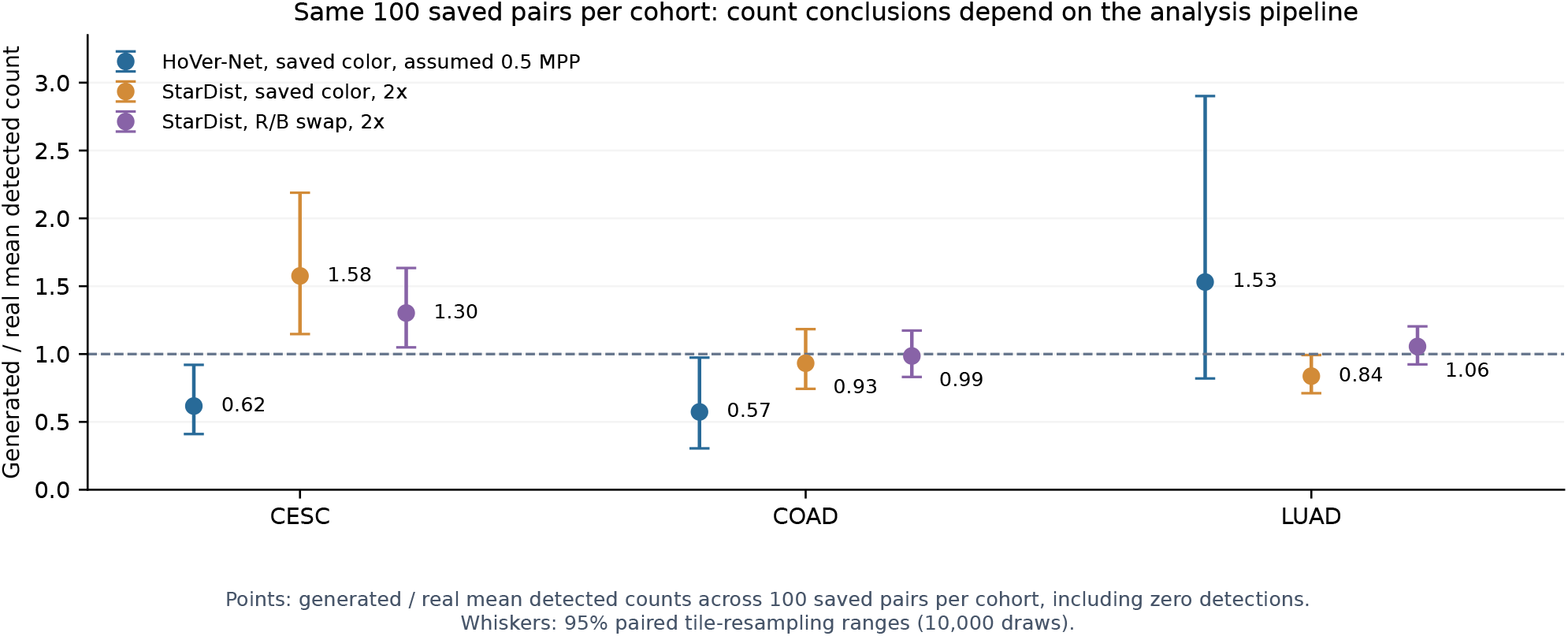
Detected-count sensitivity on identical saved indices: 100 real/generated pairs per cohort. Points show the ratio of generated to real mean instance counts; whiskers are conditional paired tile-resampling ranges. The horizontal line marks equal mean detection. Historical HoVer-Net assumes 0.5 MPP; StarDist uses 2× resampling with saved or R/B-exchanged color.

**Figure 9.**
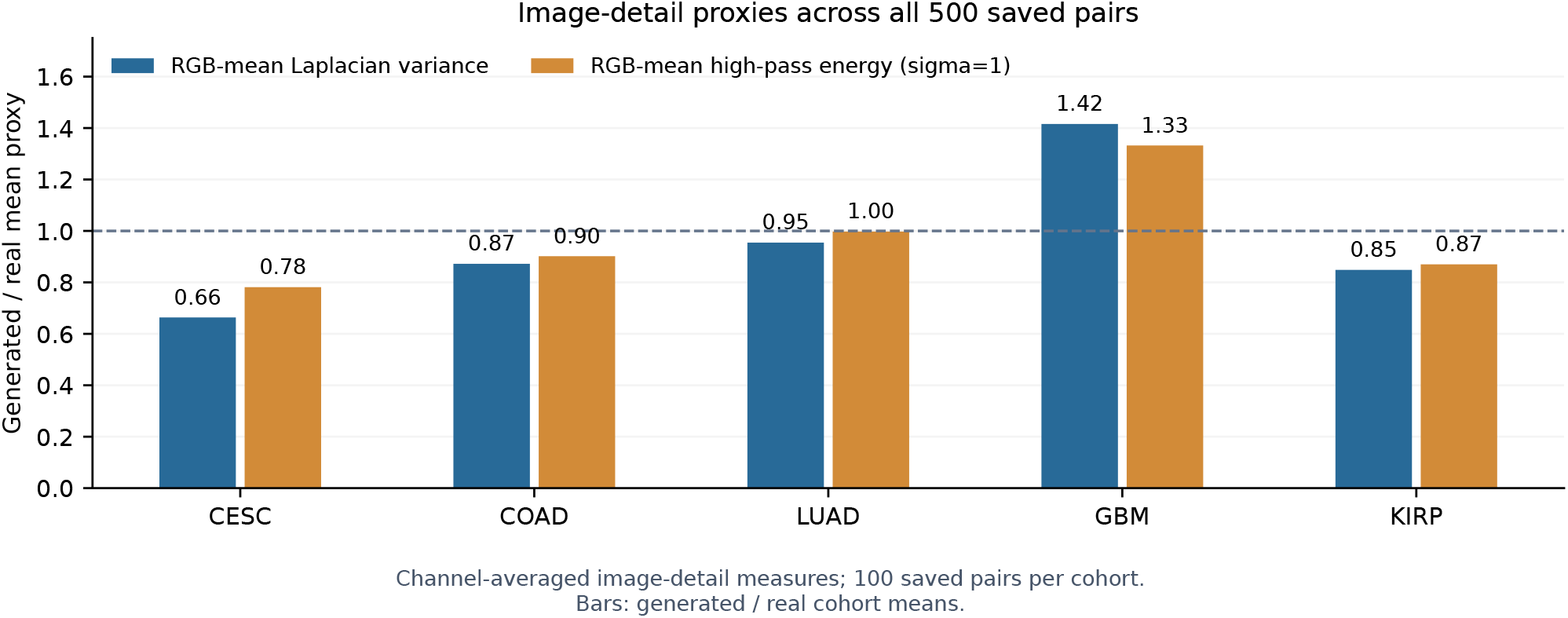
Channel-permutation-invariant image-detail proxies over all 500 saved pairs. Each proxy is computed per RGB channel and then averaged: Laplacian variance and high-pass residual energy with Gaussian *σ* = 1. Bars divide generated by real cohort means, using all 100 pairs per cohort. These measures summarize image intensity variation and structure.

#### Detector response and image detail

Ten real-image legacy outputs reproduce exactly. Five independent stitching checks reproduce all three raw output heads, and final instance counts equal retained postprocessing markers in 40 checked conditions. The tested stitching and export paths thus preserve detector outputs. In LUAD real tile 95, R/B exchange changes the raw nuclear-foreground fraction above probability 0.5 from 0.072% to 19.95% before contour counting. This localizes a measurable input response to the detector itself.

Image detail also affects detector response. Across 100 CESC pairs, channel-averaged Laplacian variance has a generated/real mean ratio of 0.664, with 85 generated tiles below their saved partners (Appendix C). On the public RGB control, Gaussian blur with *σ* = 2 changes HoVer-Net recall from 0.779 to 0.637; fixed-reference Reinhard normalization and the JPEG control give recalls of 0.743 and 0.759. Laplacian variance measures intensity variation across tissue structures. These controls quantify detector responses to changes in image detail, stain normalization, and encoding.

## 5 Conclusion

We presented RNA-SpecFlow, a continuous-latent FlowAR architecture with repeated RNA conditioning at adjacent scale transitions. A shared spectral module applies independent molecular gains to low- and high-frequency features, while a latent adapter supplies the RNA embedding and an auxiliary objective encourages its retention in backbone features. On public annotated images, twofold enlargement increases detected-nucleus counts while reducing instance agreement. On saved CESC and LUAD real/generated pairs, changing the analysis pipeline reverses the direction of detected-count differences.

Future experiments will compare matched architectural variants and RNA interventions on patient-disjoint data.

## A Display selection and artifact traceability

The fixed saved indices in Table 2 appear in both appearance and contour figures. The selection manifest records the four generated-side image proxies and the deterministic A/B rule. Nuclear predictions are not used to choose displayed tiles. Selection is deterministic and follows the saved display indices.

**Table 2.** Saved composite indices used for all A/B tissue and contour panels.

| Sample | LUAD | GBM | KIRP | CESC | COAD |
| --- | --- | --- | --- | --- | --- |
| A | 68 | 70 | 23 | 74 | 23 |
| B | 47 | 27 | 85 | 82 | 82 |

All source composites have 512 horizontal by 256 vertical pixels. Lossless PNG halves used by the nucleus analyzer are derived from these JPEGs; the halves retain the JPEG-decoded pixels. Input manifests record source hashes and crop coordinates, and instance JSONs record weight hashes, scale assumptions, probabilities, and native-pixel contours. Contours use the model-resolution-to-native factor 0.25*/*0.5 in the main analysis and factor one in the 0.25-MPP pilot. This factor maps contour coordinates to the native image grid.

## B Analyzer coverage and resolution diagnostics

Table 3 includes all 100 tiles on each side per cohort, with zero-instance tiles retained in medians and totals. A tile can have detected instances but undefined composition if all instances are Unclassified; this explains why zero-instance and undefined-composition counts differ. Marginal distributions use side-specific eligible tiles; paired differences require eligibility on both sides.

**Table 3.**
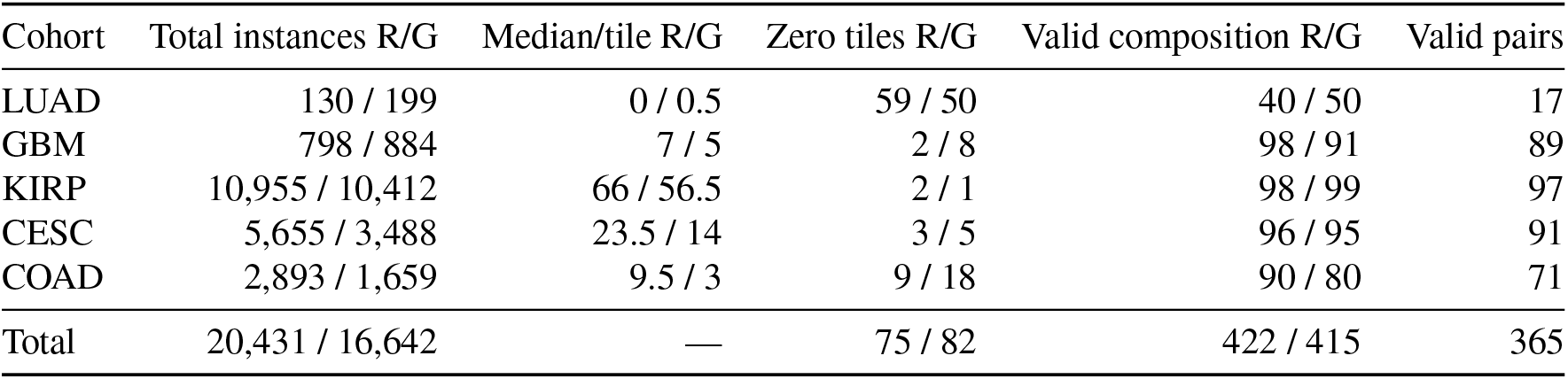
Fixed-analyzer diagnostics under the operational 0.5-MPP input assumption. R/G denotes real/generated. Each cohort has 100 saved pairs.

Table 5 gives the plotted macro estimates. Percentile ranges describe saved-index resampling under the analyzer protocol.

**Table 4.** Resolution sensitivity on the fixed Sample A tiles. Counts are real/generated predictions with identical model weights.

| Cohort | Assumed source MPP 0.5 | Assumed source MPP 0.25 |
| --- | --- | --- |
| LUAD | 0 / 2 | 0 / 0 |
| GBM | 4 / 5 | 0 / 0 |
| KIRP | 53 / 46 | 0 / 3 |
| CESC | 45 / 16 | 1 / 1 |
| COAD | 11 / 2 | 1 / 0 |
| Total | 113 / 71 | 2 / 4 |

**Table 5.** Equal-cohort paired analyzer shifts in percentage points. All rows use 365 complete pairs; each cohort receives one-fifth weight regardless of eligible-pair count.

| Predicted class | Generated – real | 95% tile-resampling range |
| --- | --- | --- |
| Tumor | +4.194 | [−2.029, 10.666] |
| Inflammatory | −5.943 | [−11.061, −0.981] |
| Connective | −1.533 | [−5.596, 2.441] |
| Dead | +0.927 | [−4.379, 6.413] |
| Normal | +2.355 | [0.792, 3.949] |

## C Expanded calibration protocol and detail proxies

The public calibration images come from the RationAI/PanNuke data mirror, fold 2. The fixed data revision is:

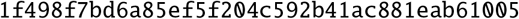

Selection takes the first three eligible rows per tissue with at least ten disjoint masks. Cervix row 970 is excluded for two overlapping mask pixels before prediction. Rows and retained instance counts are Colon 677/24, 678/13, 681/18; Lung 771/22, 772/38, 773/23; Kidney 798/24, 799/25, 800/19; and Cervix 969/33, 971/42, 1171/22. The total is 303 instances. Classes are merged for instance matching.

For each image, let TP, FP, and FN denote one-to-one matches, unmatched predictions, and unmatched annotations at IoU *>* 0.5. Then 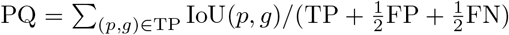. Reported PQ averages the 12 image values; precision and recall pool counts over images. An independent direct-IoU audit of 168 prediction masks agrees with the reported TP and native instance counts. Boundary fragments are retained on both sides.

HoVer-Net’s native RGB condition has 236 TP, 51 FP, and 67 FN; twofold enlargement has 197 TP, 218 FP, and 106 FN. Native R/B exchange has 41 TP, 41 FP, and 262 FN. Native RGB StarDist has 190 TP, 34 FP, and 113 FN. Twofold enlargement resamples the input pixels. Gaussian blur uses *σ* = 1 or 2 native pixels. Reinhard normalization uses a preselected Colon row 677 reference. The JPEG control uses the historical PIL quantization tables and 4:2:0 chroma subsampling.

StarDist inference uses the released 2D_versatile_he weights, probability threshold 0.692478, NMS threshold 0.3, and channelwise 1st/99.8th percentile normalization. Native/2× labels denote pixel resampling. Alternate HoVer-Net MoNuSAC weights and fixed-reference stain normalization are also examined on selected historical tiles. Results are grouped by weight set and its class definitions.

Hashes identify all 500 composite JPEGs, 1,000 source PNG halves, and legacy composition/contour figures. Transformed diagnostic inputs are stored separately. Analysis uses the readable root-level evaluation archive. Separate manifests record subset selection, inference, scoring, resampling, and figure sources.

## D Analysis reproducibility

Nuclear analysis uses the recorded detector weights, environments, source images, and preprocessing settings. Hashes identify input files and model weights. The source-scale pilot covers ten tiles. Separate manifests record subset selection, inference, scoring, resampling, and figure sources.

